# Cortical Adaptation in the Delta Band in People with Post-Stroke Aphasia during Picture Naming

**DOI:** 10.1101/2023.07.04.544349

**Authors:** Malo Renaud-D’Ambra, Alexandre Aksenov, Quentin Mesnildrey, Gesa Hartwigsen, Vitaly Volpert, Anne Beuter

**Affiliations:** CorStim SAS, Montpellier, France; Lise Meitner Research Group, Cognition and Plasticity, Max Planck Institute for Human Cognitive and Brain Sciences, Leipzig, Germany; Institute Camille Jordan, UMR 5208 CNRS, Lyon 1 University, Villeurbanne, France

**Keywords:** Language, Stroke, Aphasia, EEG, Delta band, ERP, Cortical dynamics

## Abstract

This study explores differences in spatiotemporal cortical dynamics between five people with chronic post-stroke aphasia and five healthy subjects. Electroencephalography was recorded during picture naming in both groups.

Frequency-specific Global Field Power (GFP) results showed that the delta band has higher power to discriminate between healthy subjects and people with aphasia (PWA) than theta and alpha bands. EEG topologies computed at the time of GFP peaks in the delta band revealed strong activation oscillating between posterior and anterior areas in PWA. On the other hand, EEG topologies from healthy subjects were variable.

Then, Spatial ERP (S-ERP) were developed to add spatial resolution to classical ERP analysis. S-ERP and associated analyses confirmed the previously observed oscillatory pattern in the delta frequency band among PWA during picture naming. This oscillating pattern was alternating between occipital and prefrontal areas with almost opposite phases, a characteristic not observed in healthy subjects. In addition, all PWA performed well on the picture naming task, suggesting that this oscillating pattern may be a cortical adaptation mechanism enabling them to succeed.

The observation of large-scale oscillating delta activity across the scalp in all post-stroke subjects who have substantially recovered from aphasia holds the potential to inspire innovative rehabilitation methods employing non-invasive brain stimulation techniques such as transcranial alternating current stimulation (tACS).

## 1. Introduction

Language is organized in distributed networks in the human brain. Across the last decades, our knowledge about the functional organization and specialization of language networks has considerably improved. Neuroimaging and lesion studies have identified key areas for language comprehension and speech production and point towards both overlap and specialization of networks for specific linguistic functions (Price, 2000, 2010; Vigneau *et al*., 2011; Hodgson *et al*., 2021).

Moreover, recent research provides insight into the spatial dynamics of language reorganization after brain lesions (Saur *et al*., 2006; Stockert *et al*., 2020). However, less is known about the spatiotemporal dynamics of language network in the healthy and lesioned brain. A better understanding of such processes may help to refine current models of language (re-)organization and ultimately improve current treatment approaches for people with speech and language impairments after brain lesions. One needs to clarify the link between spatial network organization and temporal dynamics during language processing.

Decades of research in the field of language production have enabled to describe the spatiotemporal cortical dynamics in finer details (Vigneau *et al*., 2006; Llorens *et al*., 2011; Liljeström *et al*., 2015; Klaus *et al*., 2019; Piai *et al*., 2019; Sarubbo *et al*., 2020; Ala-Salomäki *et al*., 2021) and to propose corresponding models of cortical processing (Strijkers *et al*., 2010; Indefrey, 2011; Laganaro, 2017). Several models of speech production exist but, despite their differences in information flow dynamics, they agree that lexical semantic information flow is accessed prior to lexical phonology during picture naming (see Graves *et al*., 2007). Accordingly, one may summarize the main steps of the picture naming process as follows: visual processing, visual recognition, semantic processing, lemma retrieval, phonological encoding, articulation programming, speech production, in parallel with self-monitoring (e.g., Indefrey and Levelt, 2004; Dell, Martin and Schwartz, 2007).

With respect to the underlying neural substrates of language, several key areas have been identified for lexical encoding (e.g., posterior middle temporal gyrus), semantic processing (e.g., angular gyrus, anterior inferior frontal gyrus), or phonological processing (e.g., inferior frontal gyrus, supramarginal gyrus), (Devlin *et al*., 1998; Vigneau *et al*., 2006; Moliadze *et al*., 2019; Stockert *et al*., 2020). However, apart from the spatial location of these areas, the mechanisms of their synchronization and/or sequential activation, their functional and/or anatomical relationship with homologue regions and neighbor brain areas remain unclear and variability among studies is important (Hampshire *et al*., 2010; Mehrkanoon *et al*., 2014; Hartwigsen *et al*., 2020, 2021). Consequently, the dynamics of such network interactions remain largely unexplored. Finally, how brain lesions affect network interactions and whether a lesion changes the cortical spatiotemporal dynamics of language production still remains unclear.

Considering such complex and high-level mechanisms, cortical damage resulting from various causes (e.g., stroke, trauma, tumor) inevitably induces physiological disorders. These disorders affect executive or cognitive capacities with different degrees of severity depending on numerous factors. Such factors include the location of the lesion, its spatial spread and depth. In response to these disturbances, complex plastic changes and reorganization are being observed either by an upregulation of network in the vicinity of the lesioned area or by the recruitment of contralateral homologous network depending on the location of the lesion (Hartwigsen, 2016; Hartwigsen et al., 2020, 2021).

The aim of this study was to identify differences in the cortical dynamics between healthy subjects and PWA. First, average GFP across subjects were investigated to assess the differences in the cortical dynamics between the two groups and a difference in the amplitude of GFP peaks in the delta band (2-4Hz) was found. Second, EEG topologies were performed at the time of the peak of GFP in the delta band and showed an oscillating pattern alternating between anterior and posterior cortical areas (Figure 4.C.G.H). To investigate this oscillating pattern a technique, which uses the spatial information, was needed. Therefore, we developed a technique of analysis based on the work of Lehman and colleagues (Lehmann *et al*., 1987), considering the scalp as a whole continuous signal using interpolation between electrodes instead of measuring each electrode independently. From this interpolation we extracted the position of maximum amplitude at each moment of time and characterized it by dividing of the scalp into 9 regions. By averaging the results on all epochs, we obtained specific signals for each region, which indicate the frequency of presence of an amplitude maximum among the 100 epochs and at each instant. As this approach is inspired from classical ERP analyses and adds a spatial component to the result, we called it Spatial-ERP (S-ERP). This new analysis revealed a new behavior (oscillating antero-posterior pattern) observed among the PWA. On the other hand, the healthy subjects presented a high variability, which impeded detecting any such pattern.

## 2. Methods

### 2.1. Subjects

Ten adult subjects took part in this study. Five of them were healthy subjects and the other five were PWA. The two groups were matched for age and gender (see details in Table 1). All subjects were right-handed, assessed using the Edinburgh Handedness Inventory (Oldfield, 1971, see scores in Table 1). PWA (labelled A*XX*) were all diagnosed with Broca’s aphasia and a moderate degree of dysarthria resulting from a first-ever ischemic stroke of the left middle cerebral artery affecting the sylvian area. They were all in chronic phase, at least six months post stroke. The exact location of the lesioned brain area as well as the severity of their language and/or motor disorders varied across subjects. A02, A04, and A05 suffered from severe motor deficits while A01 and A03 did not. The entire protocol was approved by the local ethics committee (Institutional review board, IRB-Euromov, number 2111D), all tests were conducted with the subjects written informed consent and subjects were paid for their involvement. All subjects were screened for normal cognitive functions using the Mini-Mental State Examination (score> 20, Derouesné *et al*., 1999; Kalafat *et al*., 2003). The Language Screening Test Flamand-Roze *et al*., 2011 was realized by PWA in order to provide a quick and objective measure of the type and degree of their aphasia prior to testing without diagnostic purposes.

**Table 1.** Subjects’ details. From left to right: Subject number, age at testing, sex, duration since stroke, naming score (%), Edinburgh test, screening test, cognitive test outcome measures and lesion information.

| Subject | Age (years) | sex | Duration since stroke (years) | Naming score (n=100) | Right Handedness dominance (%) | Aphasia screening test | MMSe | Stroke lesion |
| --- | --- | --- | --- | --- | --- | --- | --- | --- |
| C01 | 49 | M | n/a | 93 | 100 | n/a | 24 | n/a |
| C02 | 56 | M | n/a | 99 | 100 | n/a | 29 | n/a |
| C03 | 71 | F | n/a | 92 | 100 | n/a | 25 | n/a |
| C04 | 71 | M | n/a | 95 | 100 | n/a | 28 | n/a |
| C05 | 57 | F | n/a | 97 | 100 | n/a | 29 | n/a |
| A01 | 50 | M | 3 | 97 | 100 | 13 | 26 | Left frontoparietal<br>Deep and superficial |
| A02 | 56 | M | 4.5 | 97 | 100 | 15 | 28 | Left frontoparietal,<br>Deep and superficial |
| A03 | 70 | F | 4 | 88 | 89 | 13 | 22 | Left temporoparietal,<br>posterior sylvian area<br>Superficial |
| A04 | 71 | M | 20 | 88 | 100 | 13 | 27 | Left temporal,<br>Superficial |
| A05 | 56 | F | 3 | 89 | 100 | 15 | 26 | Left frontotemporal,<br>Superficial |

All data were acquired during a single session lasting two hours. During these two hours the subject realized the EEG, and the subject performed the task. All tests were conducted in a quiet room.

### 2.2. Picture naming

The subjects sat approximately 60 centimeters away from a computer screen. A single experimenter sat outside their visual field and provided instructions to the subject.

The main task consisted in self-paced picture naming. At the beginning of each trial, a grey screen was displayed for 1.5 seconds to allow subjects to breathe and blink naturally. Subjects were then asked to fixate a white cross displayed at the center of the screen while limiting facial movements. After 1.5s +/ − 0.2 seconds a black and white picture was displayed, and subject had to name the object at their own pace and to provide one single response. For each trial, subjects’ oral response was recorded using an external microphone located halfway between the subject and the computer screen. The voice onset was automatically detected, and a marker was added accordingly to estimate the naming latency. The next trial was automatically launched 4s after the voice onset. The coherence of automatic voice detection (thresholding) and the actual audio signal was manually verified offline and corrected when necessary to evaluate the naming latency (e.g., background noise, hesitation, or multiple responses).

Pictures were selected from the Snodgrass corpus (Alario *et al*., 1999). The set of 100 images was divided in two runs of 50 items with a small rest between them. Among the 100 images, three items were repeated 10 times each and were referred to as *control items* (“DOG”,” APPLE” and “BED”) the other 70 images were presented once. Only frequent nouns were included (*familiarity > 1*.*3*). To increase difficulty and decrease boredom, 8 out of 100 words with higher complexity were included (*familiarity* < 2, *visual complexity* > 4, see Alario & Ferrand, 1999). All items were presented in a randomized order that differed among subjects.

Four minutes resting state EEG was recorded before and after the picture naming task. For the first two minutes, subjects were instructed to look at a white fixation cross placed in the middle of the computer screen (eyes-open condition), and then close their eyes for the last two minutes (eyes-closed condition).

### 2.3. Electroencephalogram

All recordings were made using the Starstim-32 system (Neuroelectrics®). Attention was given to prevent acoustic and electromagnetic noise during the EEG acquisition. Continuous EEG recordings were obtained via 30 contacts placed on the scalp using the electrode configuration depicted in Figure 2.A at a sampling frequency of 500 Hz. Two additional electrooculography electrodes were placed on the inferior orbit and lateral canthi of the right eye to detect eye-related artifacts. The reference electrodes (CMS and DRL) were clipped on the right ear lobe. The protocol was designed using NIC2 software (Neuroelectrics, Barcelona, Spain), PsychoToolbox Brainard, 1997 and custom Matlab interfaces (Matlab 2021, the Mathworks, Natick, CA).

**Figure 1.**
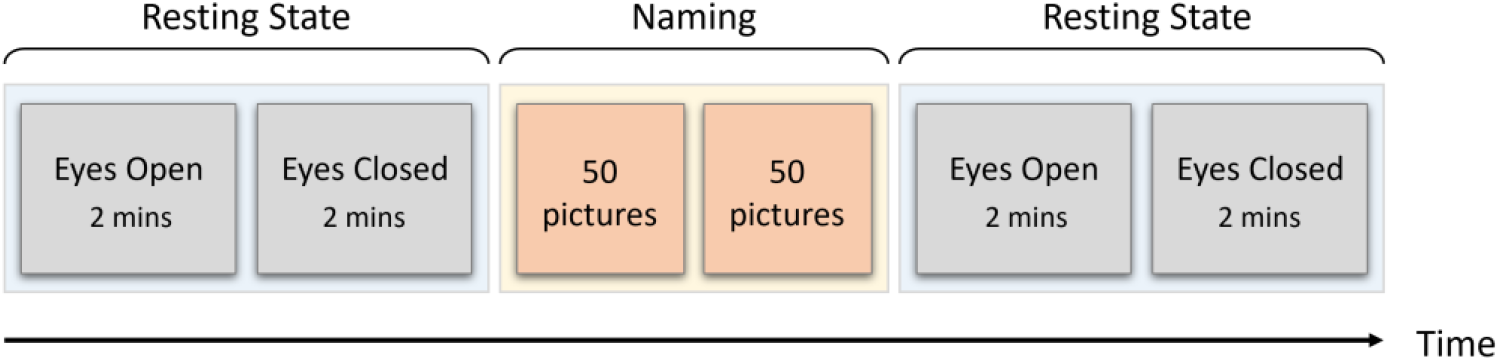
Timeline of the experimental design.

**Figure 2.**
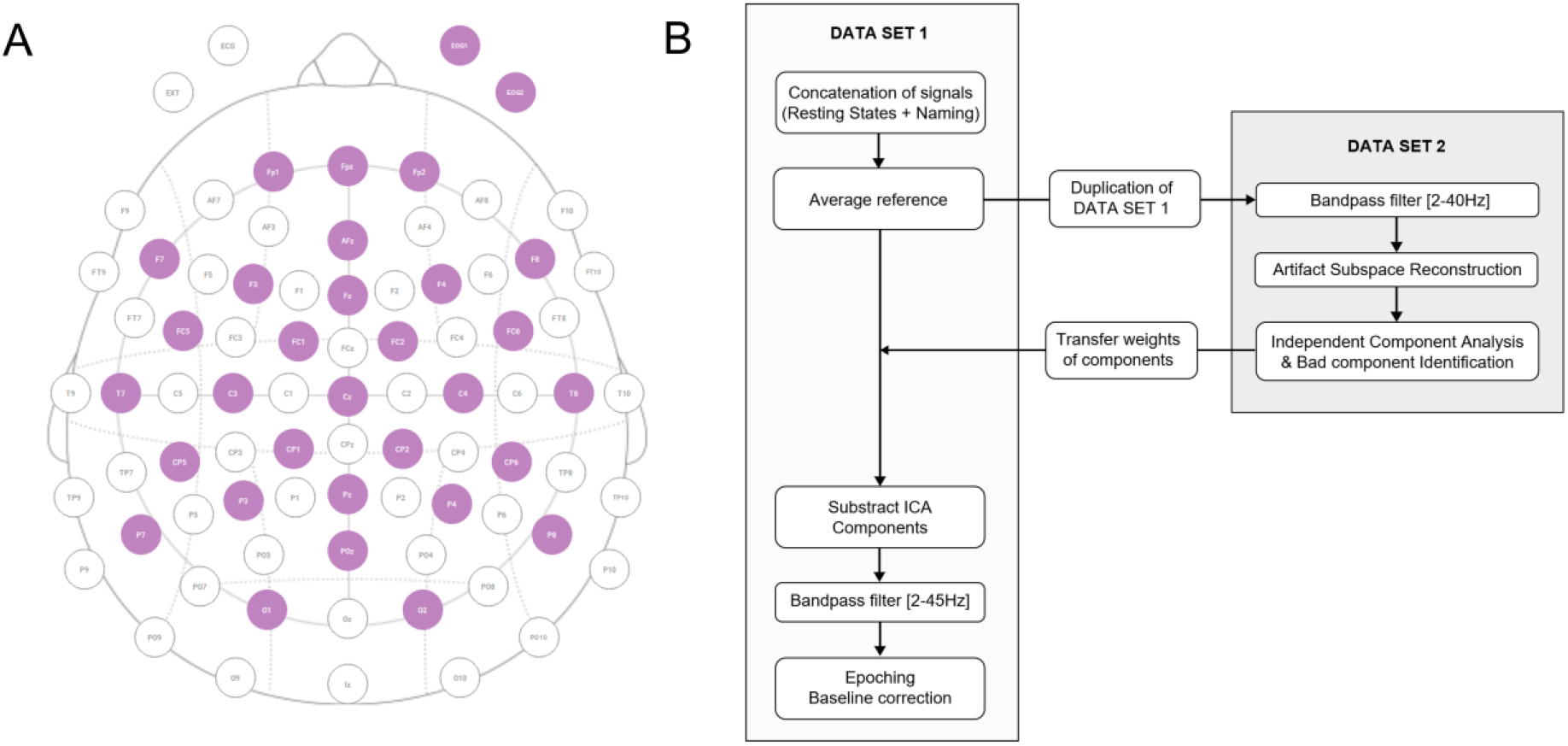
(A) Electrode configuration used for EEG recordings. Purple circles represent 30 active scalp contacts + 2 electrooculography contacts. (B) Preprocessing pipeline.

### 2.4. Analysis

#### 2.4.1. Preprocessing

Off-line signal processing and analysis were performed using EEGLAB open-source toolbox (Delorme *et al*., 2004) and custom Matlab scripts. Raw and continuous EEG signals were preprocessed using the pipeline described in Figure 2.B (EEGLAB functions). The relevance of the components of the Independent Component Analysis (ICA) was assessed, and the unwanted components were removed. Movement and muscle artifacts were also visually identified and removed when necessary. The lowest cut-off frequency of the bandpass filters was fixed at 2Hz to remove slow drifts occurring in several datasets (C01, C02, A05). The different trials (n=100) were isolated from the continuous data from 1.5s before image presentation to 4s after image presentation, thus resulting in 5.5s-long epochs. Preprocessed EEG trials were analyzed using different approaches described below.

#### 2.4.2. Global Field Power (GFP)

The global field power (GFP, Lehmann *et al*., 1980, 1984) is defined as the standard deviation across channels and thus relate to localized cortical activation or deactivation. In the present study, GFP was calculated in commonly used brain rhythm frequency ranges, such as delta (2-4Hz), theta (4-8Hz), and alpha (8-12Hz).

Signal synchronization across channels can be characterized by GFP peaks. The local peaks of the GFP were detected with a constraint preventing small peaks from being detected. Specifically, we first chose the tallest peak in each GFP signal and ignored all peaks within 50ms of it. We then repeated the procedure for the tallest remaining peak and iterated until there were no more peaks to consider. We then selected the five earliest peaks post-stimulus. The values of the GFP of the filtered signal were grouped by the frequency band, peak number, and group of subjects. The distributions of peak values of both groups of subjects were compared for each frequency band (delta, theta, alpha) and each peak number (2^nd^ to 5^th^), see Figure 3.

**Figure 3.**
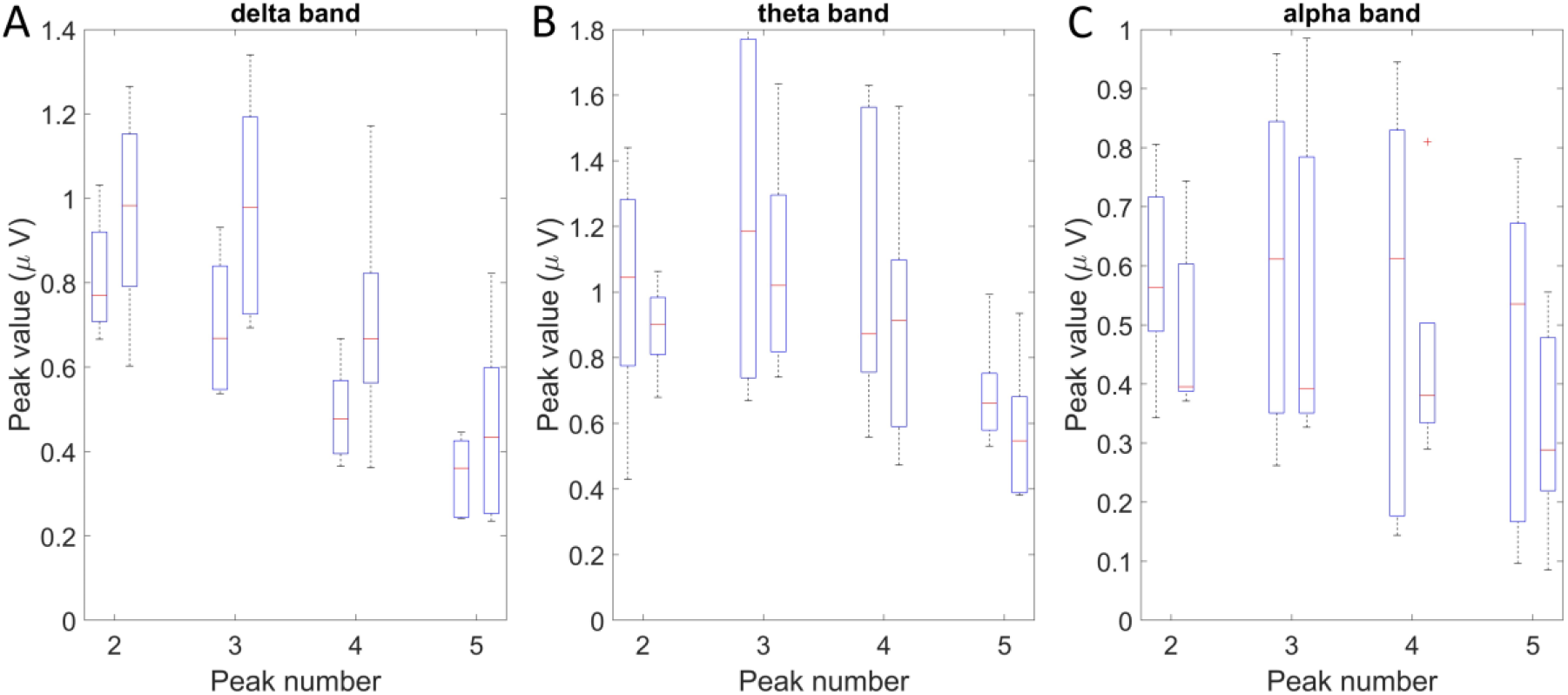
Box plot of the peak values of the second, third, fourth, and fifth peaks, in the delta band (A), theta band (B), and alpha band (C). For each peak, left box represents the peaks values of healthy subjects, right box represents the peak values of PWA.

**Figure 4.**
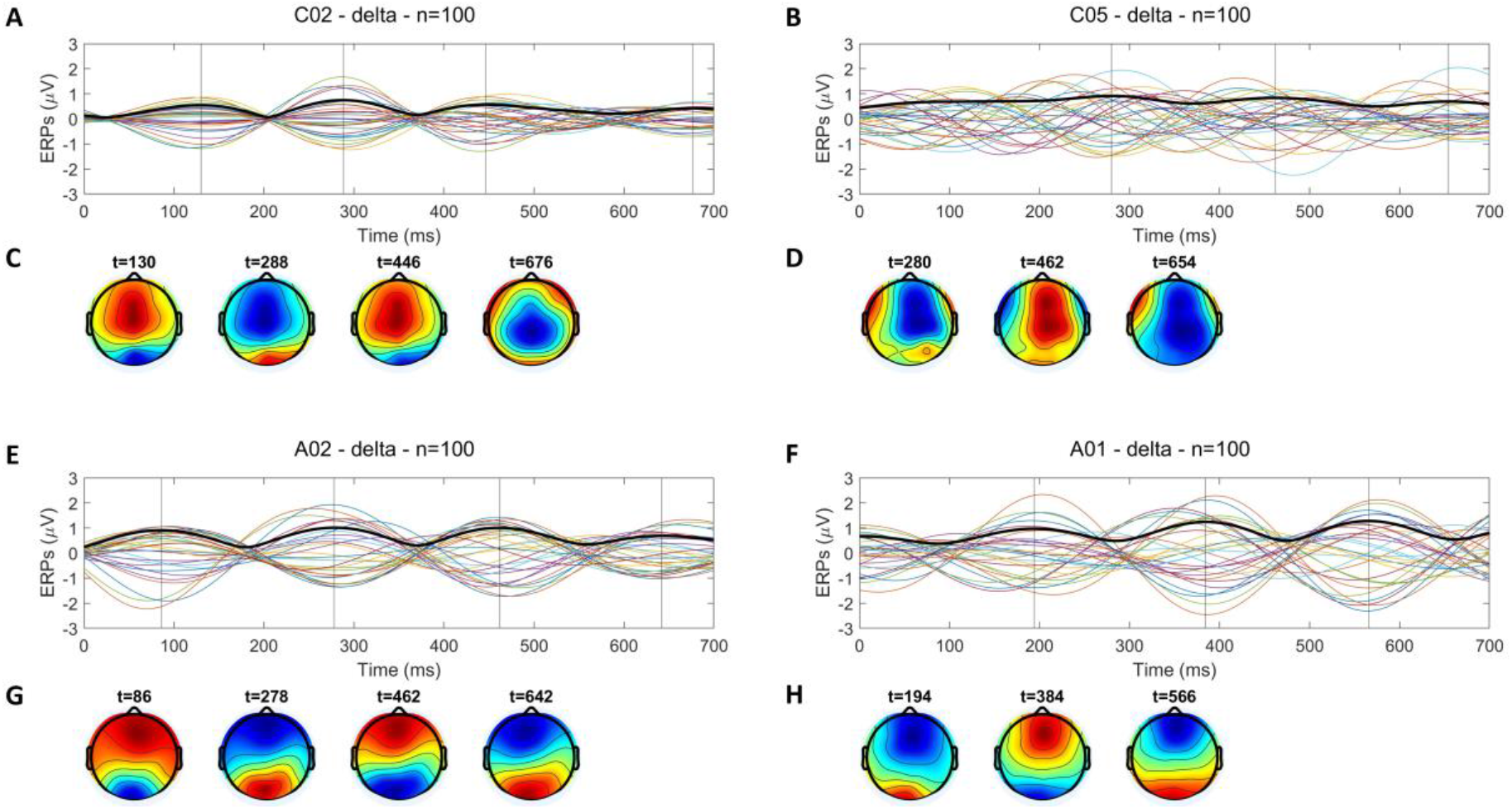
Stimulus-locked ERP signals computed from the EEG data of four subjects filtered in the delta band (2-4Hz). (A,B,E,F) ERP signals (colored lines) and global field power (thick line). (C,D,G,H) Scalp maps corresponding to the timing of GFP peaks, represented by vertical lines in panel above. (A,C) ERP signal and scalp maps of healthy subject C02. (B,D) ERP signal and scalp maps of healthy subject C05. (E,G) ERP signal and scalp maps of aphasic subject A02. (F,H) ERP signal and scalp maps of aphasic subject A01.

The first peak post-stimulus was excluded from this study for reducing the dependence of the results on the brain’s pre-stimulus activity. Indeed, the time between two peaks of bandpass-filtered GFP is expected to correspond to a half-period at the corresponding frequency (Lehmann, 1987). Therefore, the amplitude of each peak of bandpass-filtered GFP takes the brain activity during a previous of time before it into account.

The distributions of peak values were compared between groups of subjects (healthy subjects versus PWA) for each frequency and peak number. For this, a two-tailed unpaired t-test was used. The corresponding p-values quantified the amount of difference between the two groups. They were compared across frequency bands.

This analysis allowed estimating the most relevant frequency band(s) for searching differences in the cortical dynamics between the two groups as the band(s) leading the largest differences of peak values between groups. As the differences of peak values between groups were largest (measured the smallest p-values) in the delta band for each peak number from 2^nd^ to 5^th^ (see Section 3.2), this band was chosen for applying the techniques of spatiotemporal analysis.

#### 2.4.3. Event-Related Potentials (ERP) and EEG topologies

For each EEG channel, Event-Related Potentials (ERPs) were calculated as the across-trials average. As a result, the amplitude of ERP signals provides information on the repeatability across trials. ERPs are used to investigate main trends relative to a specific event (typically the appearance or change in stimulus, see Murray *et al*., 2008, for a review) with a view of assessing the correlation between an objective measure and psychophysical and behavioral observations. In the following, stimulus-locked (image presentation) ERPs were analyzed. The pre-processed EEG trials were filtered in the delta band (2-4Hz), and the duration of ERPs was 700ms.

For each ERP, the GFP of the filtered signal was also represented. Strong difference in the activity across regions is characterized by a pronounced GFP peak. The specific spatial patterns of activation are most likely to differ at the moments of pronounced peak of GFP. Therefore, EEG topologies were represented at the time points of local peaks of GFP to visualize the different patterns. EEG topologies resulted from the voltage value of the interpolated signal from all electrodes using *topoplot* function (Delorme *et al*., 2004).

#### 2.4.4. Spatial ERP (S-ERP)

Pre-processed EEG trials (n=100) of each subject were filtered in the delta band (2-4Hz). All the signals from the 30 electrodes were interpolated using *topoplot* function (Delorme *et al*., 2004). To investigate the spatiotemporal cortical dynamics, we designed a tool based on the detection of the maximum of amplitude across the interpolated signal. First, we parceled the scalp map in 9 regions (see Figure S1 in the supplementary materials) and saved the region which contained the position of maximum of this interpolated signal for every sample of each trial. Second, we computed across-trial frequency of occurrence of the maxima of amplitude in each of these different regions, for each time relative to the image presentation (Figure 5). High variation in the across-trial frequency of occurrence of the maximum of amplitude represents across-trial regularity. This approach was inspired from the work of Lehmann, *et al*., (1987) which used the maximum of amplitude of the interpolated EEG signal over the whole scalp to investigate the different functional states of the brain in the resting state. This approach called Spatial-ERP (S-ERP) is more robust to across-trial differences in amplitude but also to the noise in the signal than the ERP analysis. Indeed, it only considers the location of the maximum of amplitude for each sample of each trial. This representation enables to evaluate the relative spatial distribution of maximum of amplitude across trials for a specific subject at a given frequency. Moreover, S-ERP is more resilient to both inter-trial variations in amplitude and to the level of signal noise compared to classical ERP analysis due to the use of the location of maximum amplitude. This approach is intended as a complementary analysis rather than to replace ERP.

**Figure 5.**
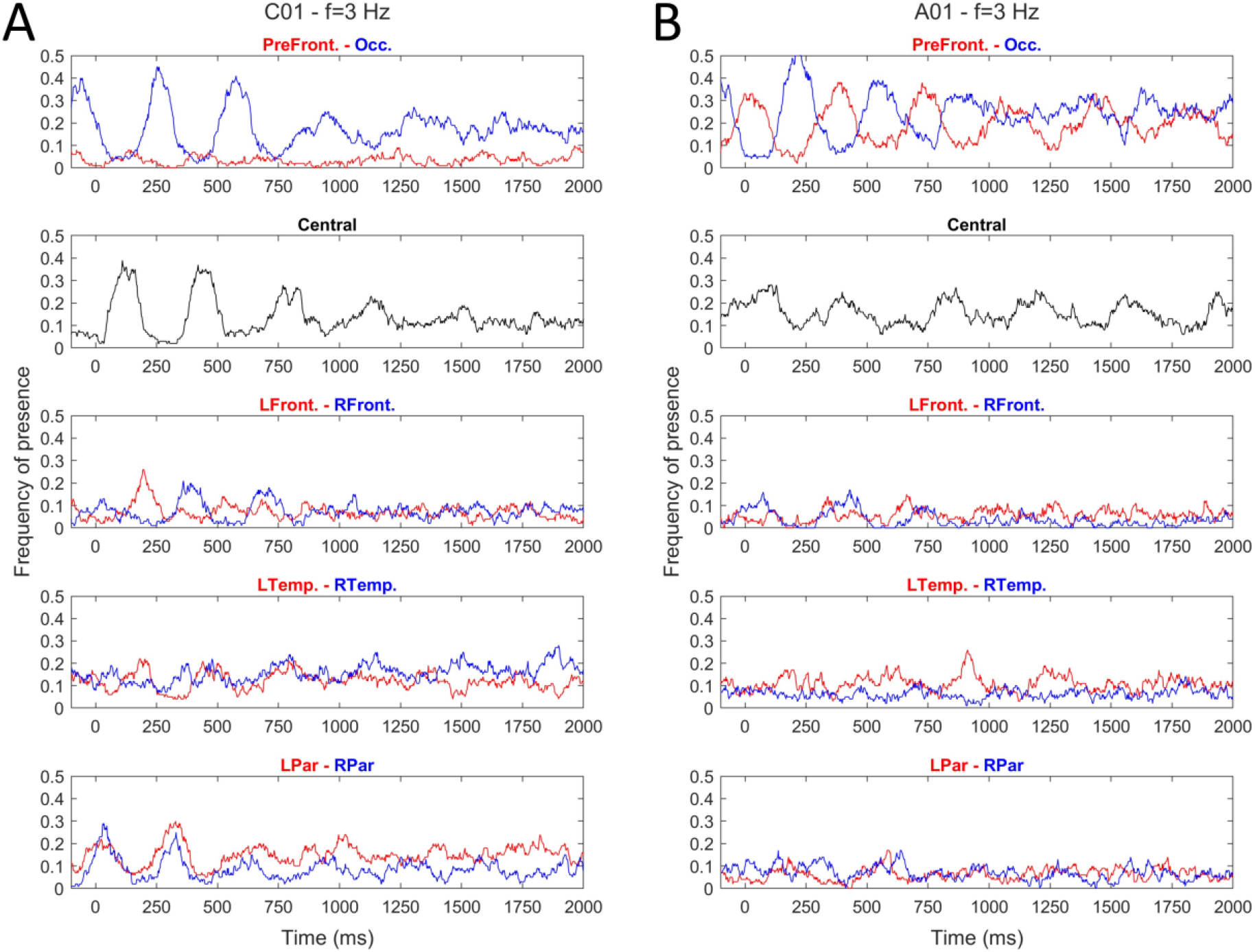
Spatial ERP showing the spatial distribution of the maxima of amplitude in the delta band. Frequency of detection of a maximum of amplitude in different scalp regions as a function of time. PreFront.=prefrontal area, Occ.=Occipital area, Central=Central area, LFront/RFront=Left/Right frontal areas, LTemp/RTemp=Left/Right Temporal areas, LPar/RPar=Left/Right parietal areas. Data from subject C01 (A), A01 (B), at a frequency of 3Hz and for all epochs, n=100.

After performing the S-ERPs, the across-trial frequency of occurrence of the maximum of amplitude, at each region for each subject, were considered as functions of time, denoted S_Region_(t). These functions were further considered as signals in time between t_1_ = -100 ms (before stimulus presentation) and t_2_ = 2000ms (after stimulus presentation) recorded at 500Hz. Techniques of signal analysis (the total power of each signal and phase angle between each pair of signals) were used to analyze their properties.

The total power of each signal S_Region_(t) was computed as its root-mean-square. The values of total power were then averaged across subjects of each group (healthy subjects or PWA).

This analysis allowed identifying the most relevant regions for searching differences in the cortical dynamics between the two groups. First, the region(s) of highest average power for each group were identified. Then, as the region(s) of highest average power was the same for both groups, the absolute value of the difference between groups of the mean power of each region was calculated, and the region with the greatest difference was identified.

#### 2.4.5. Phase angles

In addition, the phase angles were estimated between signals in all pairs of regions. For this, the cross spectral density function S_xy_(f) was estimated for all pairs of signals x = S_Region**1**_(t) and y = S_Region**2**_(t), see Section 5.2 of (Bendat *et al*., 1986). Then the phase angle *θ*_*xy*_ (*subject*) between x, y was estimated as the argument of the cross spectral density function S_xy_(f*) at the frequency f* where S_xy_(f) reached its maximum absolute value.

In this way, a phase angle was estimated at each region for each subject. The consistency of phase angles across the subjects of each group (healthy subjects or PWA) was studied using the mean resultant length associated to the phase angles (Mardia, 1972):

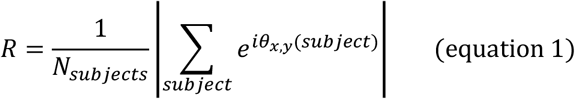

The distribution of phase angles across the subjects of each group was also tested against the null hypothesis of uniform circular distribution using the Rayleigh test. Its *p-*value was computed using the approximate formula (see Section 27 of (Zar, 2014)).

The mean phase angle:

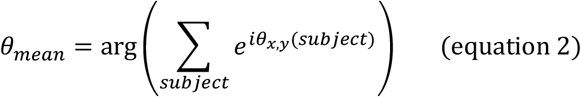

was computed as well. When the mean resultant length was high enough, the mean phase angle could be interpreted as an indication of a typical shift in time between the oscillations of activity in the two regions. For example, two regions may have a reproducible alternating behavior or a period of high activity of one region may tend to precede a period of high activity of the other.

The analysis of phase angles was performed in detail for the pair consisting of the common region of highest mean power and the region with the highest difference of mean power between groups. The resultant lengths were also computed for each group and all pairs of regions, and the resulting distributions were compared between the groups of healthy subjects and PWA.

## 3. Results

### 3.1. Behavioral results

All subjects completed the protocol. The percentage of correct naming ranges from 92 to 99% for healthy subjects and between 88 and 97% for PWA. Depending on subjects, false responses were categorized as semantic paraphasia (“*Fourchette” for” Cuillère*” i.e., “Fork” for” Spoon”), phonemic paraphasia (phoneme substitution, addition, or anticipation. e.g., “thermoter” for” thermomètre” i.e., “thermoter” for” thermometer”), failure in memory retrieval, in particular for words that are known but not frequently used. In most failed trials it was noticed that PWA were aware of providing a false response. In a few other trials, they provided a correct answer while being convinced that they failed.

The naming latency was also estimated automatically and reviewed manually when necessary. The median latency among all trials was equal to 1376ms (sd = 1114ms) for healthy subjects, and equal to 1530ms (sd = 3664ms) for PWA. The reasons for this relatively long naming latency is discussed in Section 4.1. In few occurrences, PWA were not able to provide an answer at all or provided a wrong answer after a long period of time. The naming latency could thus not be estimated for these trials. To evaluate whether PWA required a longer time for the entire process of picture naming, we compared the naming latency from successful trials only. A significant difference was found on the naming latency (for successful trials) between the two groups (*t*(933) = 3.50, *p* <0.001, mean difference = 215.5 ms). Figure S2 in supplementary materials illustrates the distribution of naming latency across groups.

Aphasia screening test scores were comprised between 13 and 15 on a total of 15. It indicated a good recovery from aphasia symptoms. MMSE were between 22 and 28 for PWA and between 24 and 29 for healthy subjects on a total of 30. It indicated that PWA had also good cognitive functions (Table 1).

In the following analyses we investigated the differences in the spatiotemporal dynamics between healthy subjects and PWA using ERP, GFP and S-ERP.

### 3.2. Global field power (GFP)

The GFP patterns showed four main peaks in both healthy subjects and PWA. The timings of these peaks were slightly delayed in PWA compared to healthy subjects. However, the delays between peak timings were relatively small and consistent within each group in the first 600ms, suggesting a synchrony of brain states across subjects and trials. It is therefore unlikely that the latency of the GFP pattern can differentiate between healthy subjects and PWA.

According to the box plot of the peak values at different frequencies (Figure 3), the peak values in theta and alpha band seemed consistent between both groups, whereas the peak values in the delta band appeared different in PWA.

Table 2 presented the p-value resulting from the two-sampled t-tests without assuming equal variances. No p-values were significant, however, p-values from delta band peaks comparisons are smaller than the ones from theta and alpha bands for all four peaks.

**Table 2.** Two-Sampled t-test. The peak values of peak number 2, 3, 4 and 5 and in delta (2-4Hz), theta (4-8Hz) and alpha band (8-12Hz) were used. T-tests were realized between peak values of healthy subjects and PWA without assuming equal variances.

|  | Delta band (2-4Hz) | Theta band (4-8Hz) | Alpha band (8-12Hz) |
| --- | --- | --- | --- |
| Peak number | p-value | p-value | p-value |
| 2 | 0.29 | 0.55 | 0.39 |
| 3 | 0.09 | 0.60 | 0.80 |
| 4 | 0.18 | 0.54 | 0.64 |
| 5 | 0.37 | 0.38 | 0.47 |

### 3.3. Event Related Potentials (ERP) and EEG topologies

The ERP and EEG topologies (see Figures 4 and S3, Supplementary Materials) showed that all PWA showed signal synchronization across channels resulting in pronounced GFP peaks, while healthy subjects showed different degrees of synchronization. EEG topologies computed at the time of pronounced GFP peaks (all PWA, and C01 and C02, see Figure S3) presented an oscillating pattern between anterior and posterior cortical areas. On the other hand, ERP signals from C03-C05 were not synchronized across channels, leading to flat GFP peaks and to EEG topologies without clear pattern.

Figure 4 displays examples of ERP signals (A,B,E,F) and the scalp maps (C,D,G,H) computed at the peaks of the GFP (illustrated by vertical lines, methods described Section 2.4.3). The scalp maps shows examples of patterns, detected at moments of pronounced (C,G,H), versus flat (D) peaks of GFP. The corresponding EEG topologies presented either an oscillating pattern, or absence of specific pattern. Additional analyses (S-ERP) were performed to quantify the potential oscillating patterns and their reproducibility across subjects.

### 3.4. Spatial ERP (S-ERP)

Figure 5 displays two examples (C01, A01) of results of the method S-ERP for EEG data filtered at the delta band. As an example, Figure 5.A shows that for subject C01, S_Occipital_(250 ms) =0.44, which means that at t=250ms post-stimulus 44% of epochs showed a maximum of amplitude in the occipital region. Another example is: S_Central_(400ms)=0.28, which means that at t=400ms post-stimulus 28% of epochs showed a maximum of amplitude in the central area.

The signal S_Occipital_(0-600 ms) of each of the 10 subjects (Figure 6 B,D) presents high variation, implying temporal regularity in the activation patterns across the 100 epochs.

**Figure 6.**
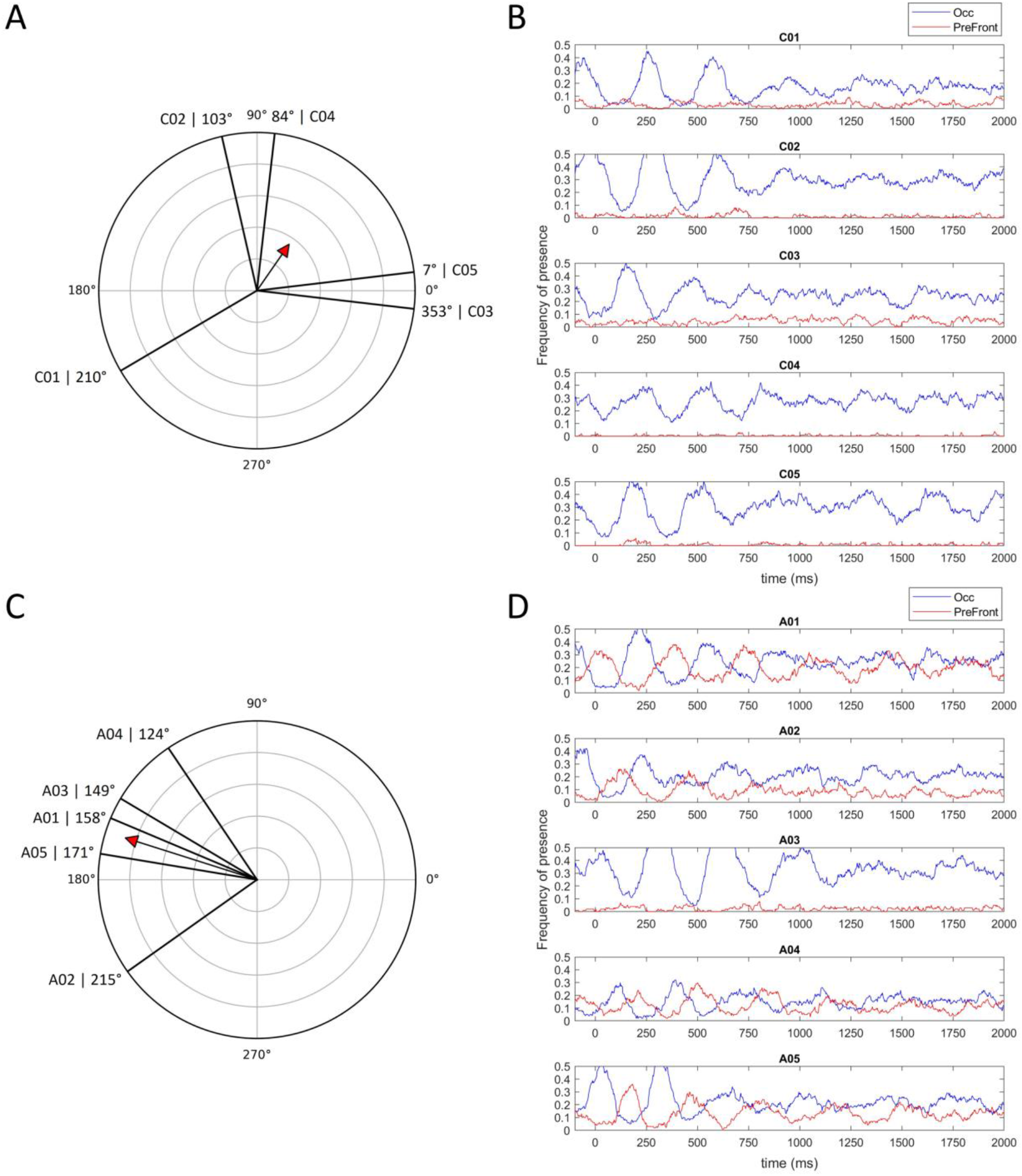
A (healthy subjects), C (PWA): rose diagrams of the phase angles estimated for the 5 subjects of each group. The plotted radii of the unit circle indicate the phase angles and the arrow indicates the mean of the associated unit vectors. B (healthy subjects), D (PWA): the S-ERP signals for the Occipital (blue) and Prefrontal (red) regions for each subject.

The region with the highest mean power across groups was the occipital region (Table 3) for both groups. The second region with the highest mean power was the central region for healthy subjects (mean power = 0.16) and for PWA (mean power = 0.13). The highest difference of mean power between groups was found in the prefrontal area. Indeed, the mean power of S_Prefrontal_(t) of healthy subjects was almost null (0.027), while PWA presented a relatively high (0.12) mean power of S_Prefrontal_(t) (Table 3). The results relative to the mean power in groups allow to study the phase angles between the regions Prefrontal-Occipital, according to the criterion Section 2.4.5.

**Table 3.**
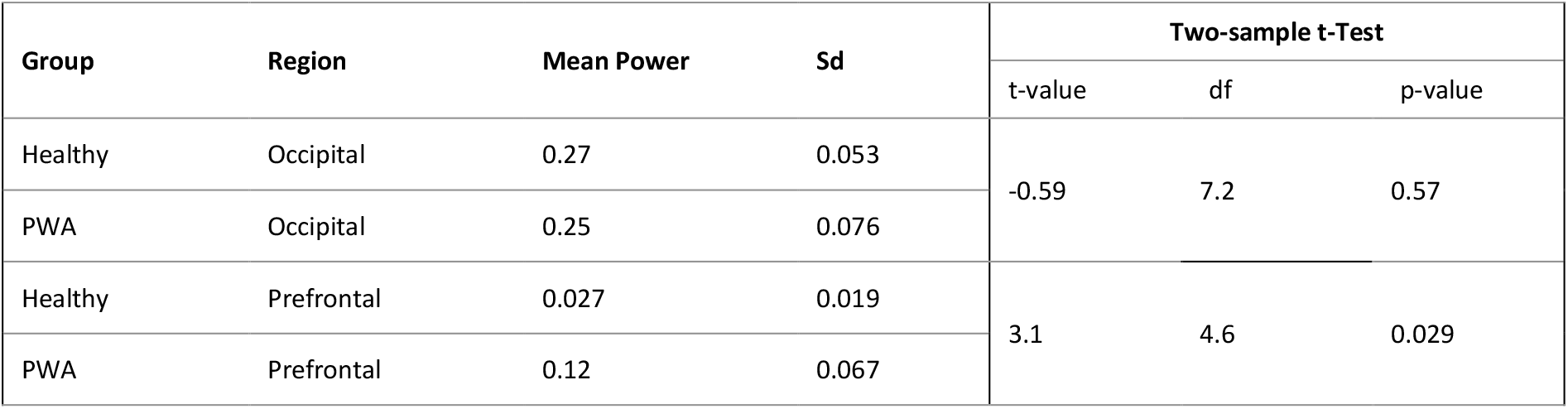
Mean power of S-ERP signals in occipital and prefrontal regions for healthy subjects and PWA. The right side of the table shows the results of two-sample t-tests on the difference in means without assuming equal variances. The Prefrontal region leads to the highest difference between groups relatively to their variance (indicated by the highest t-value is and smallest p-value).

### 3.5. Phase angles

The results concerning the phase angles (see Section 2.4.5 above) between different pairs of regions, as well as the consistency of results within each group of subjects) are presented in this section. The pair of regions Prefrontal-Occipital, selected in Section 3.4, is studied in finer detail.

Figures 6 B,D show the pairs of signals of S-ERP for prefrontal and occipital regions and all 10 subjects. This representation allows detecting their relative behavior visually: do the variations of one signal precede the variations of the other or do the signals vary in phase or in opposite phase? The answers for each are estimated using the technique of phase angles (see Section 2.4.5) and represented Figure 6 A,C. The groups are separated between panels of each figure (A,B: healthy subjects; C,D: PWA) so that the consistency of phase angles between subjects of each group can be more clearly visualized. The associated mean vectors represented Figures 6 A,C, show both the mean phase angle of the group (equation 1) and the mean resultant length (measured as the length of the mean vector, equation 2, and interpreted as a measure of consistency between the subjects of a group). The mean phase angles and mean resultant lengths for each group of subjects and pair of regions can be found Table 4.

As the mean power was quite low for the healthy subjects in the region of highest difference (prefrontal), the phase angles were analyzed as well for the two regions of highest mean power (occipital and central regions for the healthy subjects (Figure S4, supplementary materials).

The populations of phase angles for the subjects of each group were tested against the null hypothesis of uniform circular distribution using the Rayleigh test. The phase angles between occipital and prefrontal region of PWA rejected the null hypothesis (mean phase angle = 162°, R = 0.87, p-value = 0.014), while for healthy subjects these phase angles fail to reject the null hypothesis of angles selected at random (mean phase angle = 55°, R = 0.36, p-value = 0.553).

The distributions of mean resultant lengths (R) for all 36 pairs of regions for each group are represented Figure 7. The vertical lines indicate the mean resultant length R for the Occipital-Prefrontal pair for the PWA. Figure 7 shows that the Occipital-Prefrontal pair has a higher across-subject consistency (measured by the mean resultant length R) for PWA than all pairs of regions for the group of healthy subjects. Moreover, for PWA, other pairs of regions presented high consistency: occipital and left frontal (R = 0.92, mean angle = 174°), occipital and right frontal (R = 0.95, mean angle = 180°). The potential implications of this consistent behavior of PWA will be discussed Section 4.3 below.

**Figure 7.**
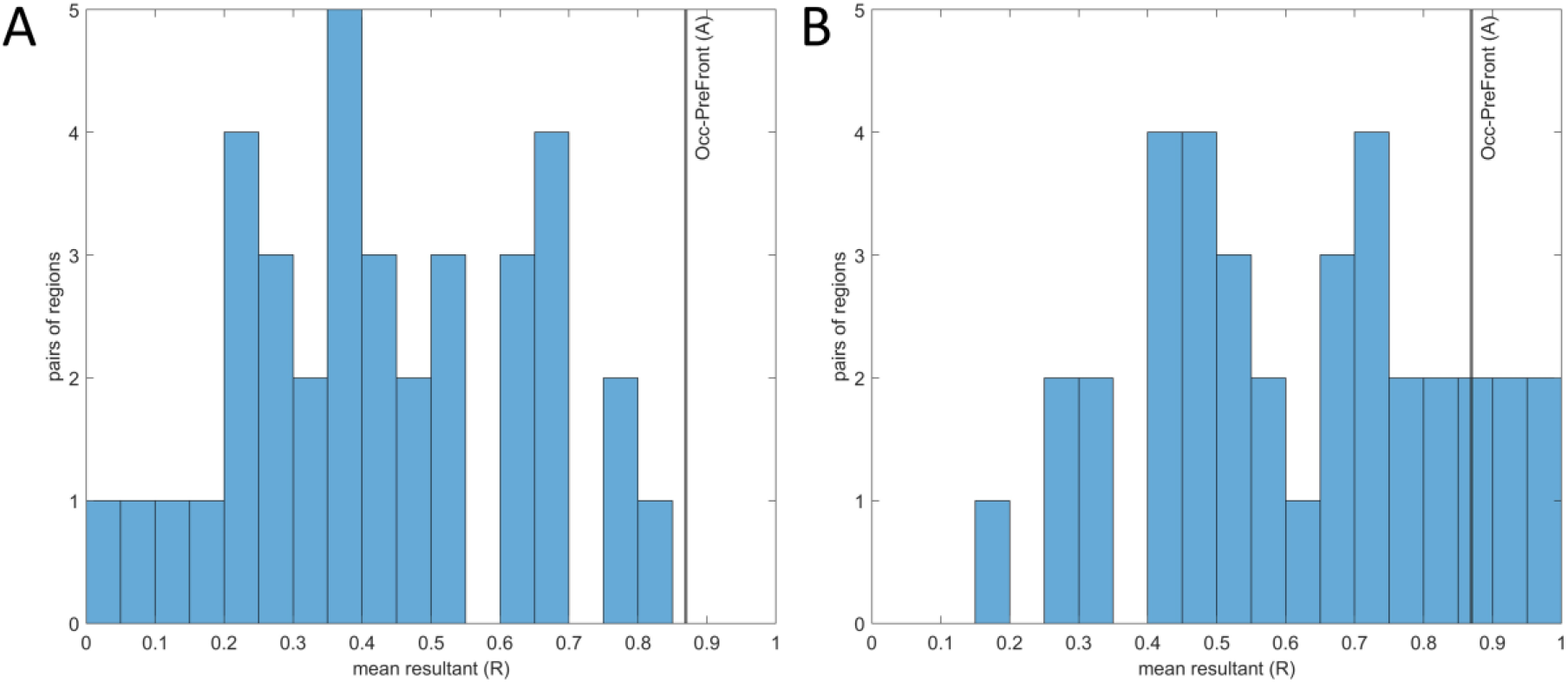
Histogram of mean resultant length (R) for all pairs of regions (each histogram represents a distribution of 36 values, which correspond to 36 pairs of regions). The vertical lines indicate the mean resultant length R for the Occipital-Prefrontal pair for PWA. A: the group of healthy subjects, B: the group of PWA.

## 4. Discussion

Previous research has established the existence of various cortical networks implicated in language production. However, the spatiotemporal dynamics monitoring the activity of these networks and how they are influenced after stroke are not well understood yet. In the present study, we investigated these dynamics by comparing cortical activity during self-paced picture naming between healthy subjects and PWA. GFP analyses highlighted differences in the delta-frequency band between both groups, so we continued the investigation in the delta band with ERPs and EEG topologies at the timing of GFP peaks. EEG topologies revealed an oscillating pattern between anterior and posterior areas, mainly in PWA. S-ERP results and associated analyses demonstrated the presence of a delta-band oscillating pattern between prefrontal and occipital regions in PWA, while healthy subjects showed high variability.

### 4.1. Behavioral correlates of language production

Behavioral results indicated that PWA perform well despite showing signs of effortful processing. Most naming errors were categorized as semantic or phonemic paraphasia. A significant difference in naming latency was observed between healthy subjects and PWA for all trials as well as for successful trials only, PWA had higher naming latency. Note that the naming latency measured in the present experiment is longer than those reported in the literature (approximately 800ms across five different studies reviewed by Laganaro, 2017, and 883ms on average for Alario & Ferrand, 1999). An explanation for the observed longer latency in our study may come from task design which allowed self-paced production without emphasizing the speed of the response, indeed the subjects were not asked to answer as quickly as possible. PWA also showed different degrees of dysarthria. However, for these subjects we consider these errors to result from a difficulty in associating phonetic and phonemic information rather than actual motor disorders (Blumstein *et al*., 1982; Kohn, 1988; Laganaro *et al*., 2010). Aphasia screening test and MMSE results underlined the good recovery in language and cognitive functions of PWA.

### 4.2. Electrophysiological correlates of picture naming

Frequency-specific Global Field Power (GFP) results showed that the delta band has higher power to discriminate between healthy subjects and PWA than theta and alpha bands. In addition, EEG topologies at GFP peaks (computed on the basis of ERP in delta band) demonstrated cortical oscillating patterns after stimulus presentation reproduced in all PWA. On the other hand, the results relative to the healthy subjects presented high variability. The cortical patterns identified in these EEG topologies reflected specific brain states (e.g., Chantsoulis *et al*., 2017; Laganaro, 2017; Mheich *et al*., 2021^)^. The observed oscillating pattern was quantified using S-ERP and phase angle techniques.

### 4.3. S-ERP revealed/highlighted oscillating pattern in delta band during picture naming

As argued by Alexander *et al*., 2013, averaging recordings such as cortical activity drastically reduces the intrinsic information. Cortical activity undergoes continuous changes, and when multiple trials are averaged, a significant portion of the information is inevitably lost. This is because averaging primarily emphasizes the contribution of phase-synchronized physiological signals, potentially overlooking other valuable aspects of the neural response. From this perspective, S-ERP analysis has the advantage of introducing spatial dimension by determining the position of the maximum amplitude of brain activity among 9 regions for every sample of each trial.

Considering the frequency of occurrence of amplitude maxima over time as a signal S_Region_(t) allowed us to apply signal analysis and to study its power (Section 3.4) and phase (Section 3.5).

#### 4.3.1. Power

Regarding power of S_Region_(t), the region with the maximum average power was the occipital region for both groups. This was consistent with the experimental paradigm since this region is involved in visual processing, and picture naming involved image recognition. The absolute difference between mean powers among groups is highest for the prefrontal region. This region was not expected to present maximal cortical activity during picture naming. Indeed, the region characterized in our segmentation as “prefrontal” includes the four electrodes “Fp1”, “Fpz”, “Fp2” and “Afz”, which does not correspond to the positions of the different areas classically involved in language (Devlin *et al*., 1998; Vigneau *et al*., 2006; Moliadze *et al*., 2019; Stockert *et al*., 2020). This result suggests that a high activity of the prefrontal region is needed for PWA to achieve picture naming, whereas it is not for healthy subjects. As an important caveat, this observation does not mean that the prefrontal cortex is not active in healthy subjects but rather that the cortical activity in the delta band was consistently larger in other regions, i.e occipital region, central region. Therefore, the presence of such high activity in this region in PWA revealed the need for strong activation of other anatomical regions than those reported in the literature as involved in picture naming in healthy subjects (Devlin *et al*., 1998; Vigneau *et al*., 2006; Moliadze *et al*., 2019; Stockert *et al*., 2020). In addition, the mean power of central region of both groups was the second higher region behind occipital region. This might indicate the early involvement of premotor areas (Llorens *et al*., 2011).

#### 4.3.2. Phase angle

In addition to having higher activity in the prefrontal region, the alternation between the occipital and prefrontal regions of PWA was almost characterized by opposite phases with a significant consistency (R) between subjects (Figure 6), a behavior that is not observed in healthy subjects. We could note that the low mean power of the signal in the prefrontal region S_Prefront_(0-1000) of healthy subjects negatively impacted the measure of its phase angle, however the distribution of measures of consistency (R) for all pairs of regions (Figure 7) revealed, for healthy subjects, no pair of regions presented such a high consistency as the one from the prefrontal-occipital pair of PWA. This result emphasized the importance of this specific oscillating pattern observed in all PWA. The other high consistencies observed between occipital and both left frontal, and right frontal regions for PWA confirmed the specific alternation of high cortical activity between anterior and posterior areas observed in EEG topologies (Figure 4).

#### 4.3.3. Delta band

Delta oscillations are regularly observed during sleep and general anesthesia, leading to the assumption that they have little relevance to cognition (Hobson *et al*., 2002, Schomer *et al*., 2011) and limited research are available on the functional significance of the delta band to date. However, Gross *et al*. (2013) demonstrated the implication of delta oscillations in language processing, specifically in the auditory cortex where speech segmentation and coding are based on a nested hierarchy of entrained cortical oscillations. Moreover, the involvement of delta activity in naming and thus in language production as well as cognitive control has been observed recently by Liu *et al*. (2022). In our study, the oscillating pattern in delta band could be implicated in a phenomenon of frequency coupling needed for the realization of picture naming. Finally, Nácher *et al*. (2003) showed that during decision making, delta-frequency band links distant cortical circuits. All these studies lead us to consider the potential involvement of this large-scale oscillatory pattern in the delta band as being effectively involved in the realization of picture naming.

Among the physiological disorders reported after a cortical injury, delta rhythm is known to be associated with brain trauma such as tumor (Jongh *et al*., 2003) and stroke (Spironelli *et al*., 2019). Indeed, several studies have reported altered cortical dynamics in the delta frequency band in PWA. In a study by Spironelli *et al*. (2009), after identifying significantly higher delta amplitude in PWA compared to healthy subjects, particularly in left perilesional EEG sites, the authors concluded that delta activity might represent a valuable and sensitive means of evaluating the reorganization and recuperation of impaired neural networks. In Meinzer *et al*. (2004) the magnitude of change of delta activity in the perilesional area of PWA was assessed before and after intensive speech and language therapy and reported both patient-specific increase and decrease in this activity (16 and 12 patients respectively). In a study examining functional connectivity, Kawano *et al*. (2021) recently developed a new approach based on the electroencephalographic phase synchrony index. They reported an increase in functional connectivity between the right frontotemporal lobes of PWA and propose it as a biomarker of post-stroke aphasia. These studies emphasized the potential implication of delta-frequency band in adaptation mechanisms.

#### 4.3.4. Adaptive mechanism

In relation to maximum amplitude, cortical mechanisms among healthy subjects exhibited substantial variation, whereas PWA displayed a more uniform pattern, suggesting the presence of an adaptation mechanism subsequent to a stroke. In this study, we intentionally included individuals with post-stroke aphasia who had experienced a left sylvian region stroke more than 6 months prior and had successfully regained proficient speech abilities. Concurrent with favorable naming scores, the observed cortical dynamics in this study provide support for the hypothesis of an adaptive phenomenon, rather than a direct consequence of the stroke.

In this study the PWA are in chronic phase (time from stroke > 6 months) and have globally well performed the task (score from 88% to 97%), thus it seems they have already developed new cortical networks to compensate for the alteration of the stroke in networks involved in language initially. Moreover, in terms of amplitude maxima, healthy subjects showed varied cortical mechanisms, while PWA showed similar ones. Therefore, considering all the clues about the implication of delta band in cognitive task and its modulation after stroke, we make the hypothesis that these specific large scale cortical dynamics in the delta band between posterior (occipital) and anterior (prefrontal and frontal) regions could be one mechanism of adaptation of PWA to achieve picture naming after stroke.

#### 4.3.5. Opening

This result adds further clues to the involvement of the delta rhythm in complex cognitive processes. Furthermore, it would be interesting to study the influence of delta rhythms in the rehabilitation of post-stroke patients, especially in language. By consolidating research on this subject, it could be interesting to investigate whether the same delta activity is observed in post stroke patients within the acute phase instead of chronic phase, it could help to define whether this oscillating pattern is due to the lesion itself, or if it is an adaptive mechanism contributing to language recovery. Moreover, non-invasive brain stimulation studies using tACS to modulate delta activity in patients could be considered as a rehabilitation method.

### 4.4. Limitations

These preliminary results are limited by the small sample size and the relatively large number of outcome variables. Consequently, our findings should be interpreted with caution. PWA had an ischemic stroke of the left middle cerebral artery affecting the sylvian area on the left hemisphere associated with mild aphasia which may have minimized the intersubject variability in cortical dynamics among the PWA. This aspect is both an advantage as it allows a more precise understanding of this specific pathology, and a disadvantage, as it does not allow generalization to all patients with post-stroke aphasia. Repeating this study with a larger cohort of patients could allow us to verify these results.

### 4.5. Conclusion

This exploratory study highlighted the involvement of the delta band in picture naming in PWA. Our results provide new clues suggesting the involvement of large-scale delta rhythm in the realization of cognitive tasks. This study also brings new evidence about the consequences of stroke on cortical dynamics, specifically in the delta band. Furthermore, we introduced a novel analysis technique called Spatial-ERP, which specifically targets the distribution of maximum amplitudes across the entire scalp, averaged across epochs. This innovative approach enabled us to identify a distinct oscillatory pattern in the delta frequency band among individuals with post-stroke aphasia during picture naming. This oscillating pattern was alternating between prefrontal and occipital areas with almost opposite phases, a characteristic not observed in healthy subjects. Associated with a high level of success in picture naming in chronic stroke patients, this cortical behavior might reveal an adaptive mechanism. Further studies are needed to confirm the results of this exploratory study which could lead to the development of new rehabilitation therapies using noninvasive transcranial stimulation to reinforce this adaptive mechanism.

## Acknowledgments

The authors sincerely thank Gérard Dray for his support in this study, all subjects of the naming experiment and the Fédération Nationale des Aphasiques de France (FNAF). This research did not receive any specific grant from funding agencies in the public, commercial, or not-for-profit sectors.

## Authors contributions

All authors were implicated in the conception, the recording, and the analyses of the experiment.

## Conflict of interest disclosure

The authors declare that they comply with the PCI rule of having no financial conflicts of interest in relation to the content of the article.

## Notes

### Competing Interest Statement

The authors have declared no competing interest.

