## Supplementary Materials for "Cortical Adaptation in the Delta Band in People with Post-Stroke Aphasia during Picture Naming"

Figure S1 : Schematic view of the parcellation of scalp surface used for Spatial ERP (S-ERP, Figure 5).

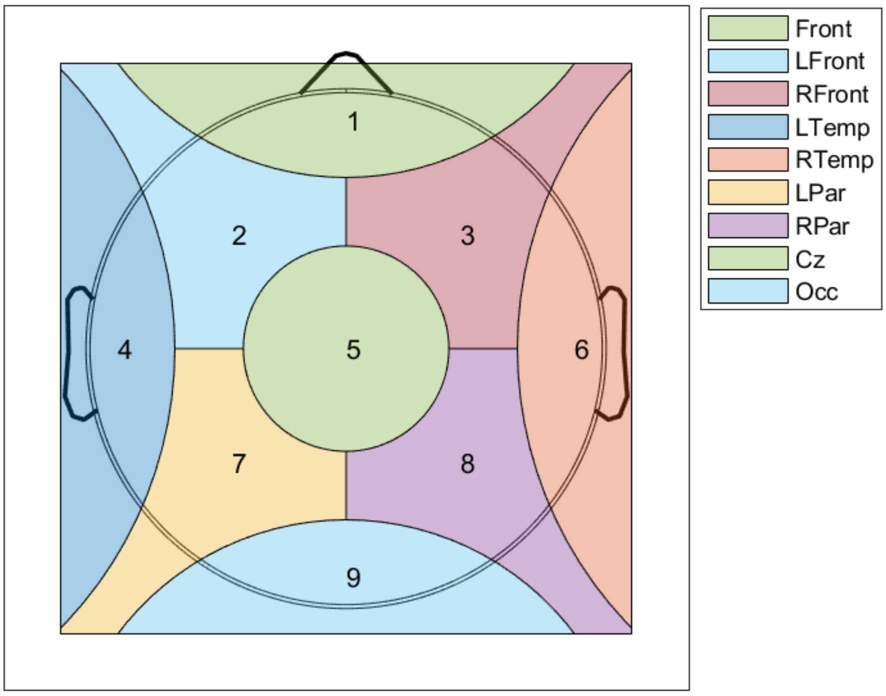

Figure S2 : Naming latency (in ms) in the group of people with aphasia (PWA) and in the group of healthy subjects. Red lines indicate the median latency, black lines indicate the mean latency.

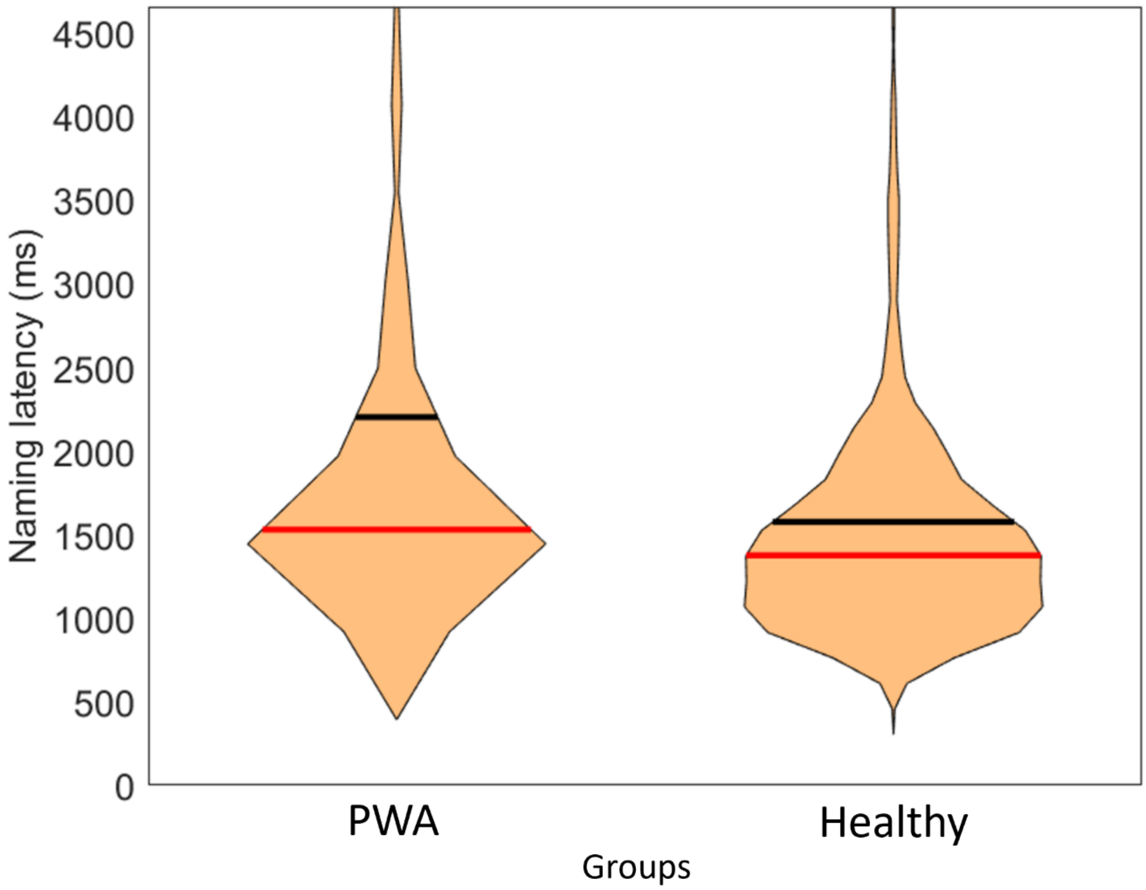

Figure S3. Stimulus-locked ERP signals computed from the EEG data of the 5 subjects of each group (C01-5: Healthy, A01-5: PWA). filtered in the delta band (2-4Hz). For each subject, ERP signals (colored lines) and global field power (thick line) are presented on the top panel. For each subject, scalp maps at the timing of Global Field Power (GFP) peaks (represented by vertical lines in the top panel) are presented on the bottom panel.

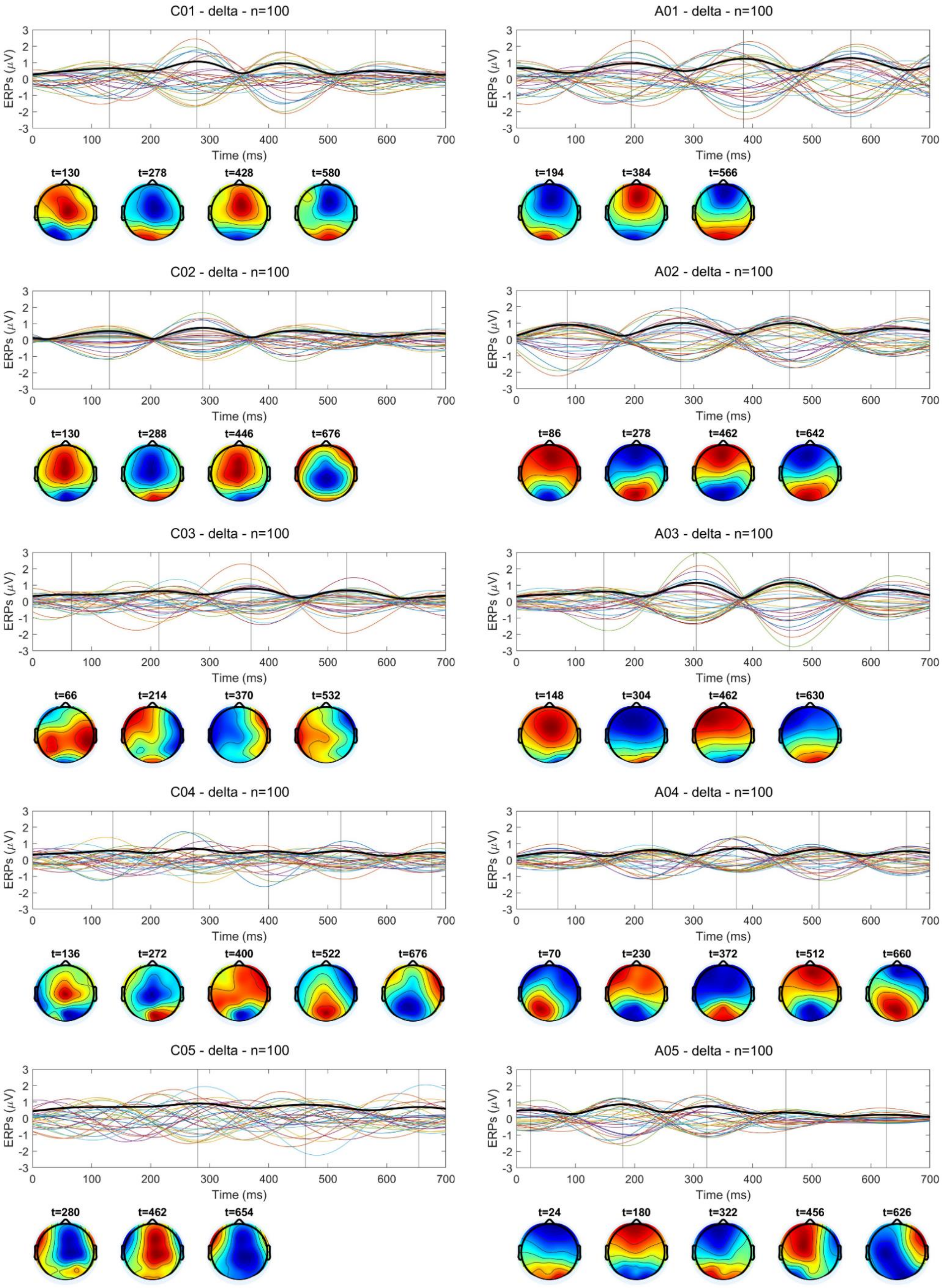

Figure S4. A (healthy subjects), C (PWA): rose diagrams of the phase angles estimated for the 5 subjects of each group. The plotted radii of the unit circle indicate the phase angles and the arrow indicates the mean of the associated unit vectors. B (healthy subjects), D (PWA): the S-ERP signals for the Occipital (blue) and Central (red) regions for each subject.

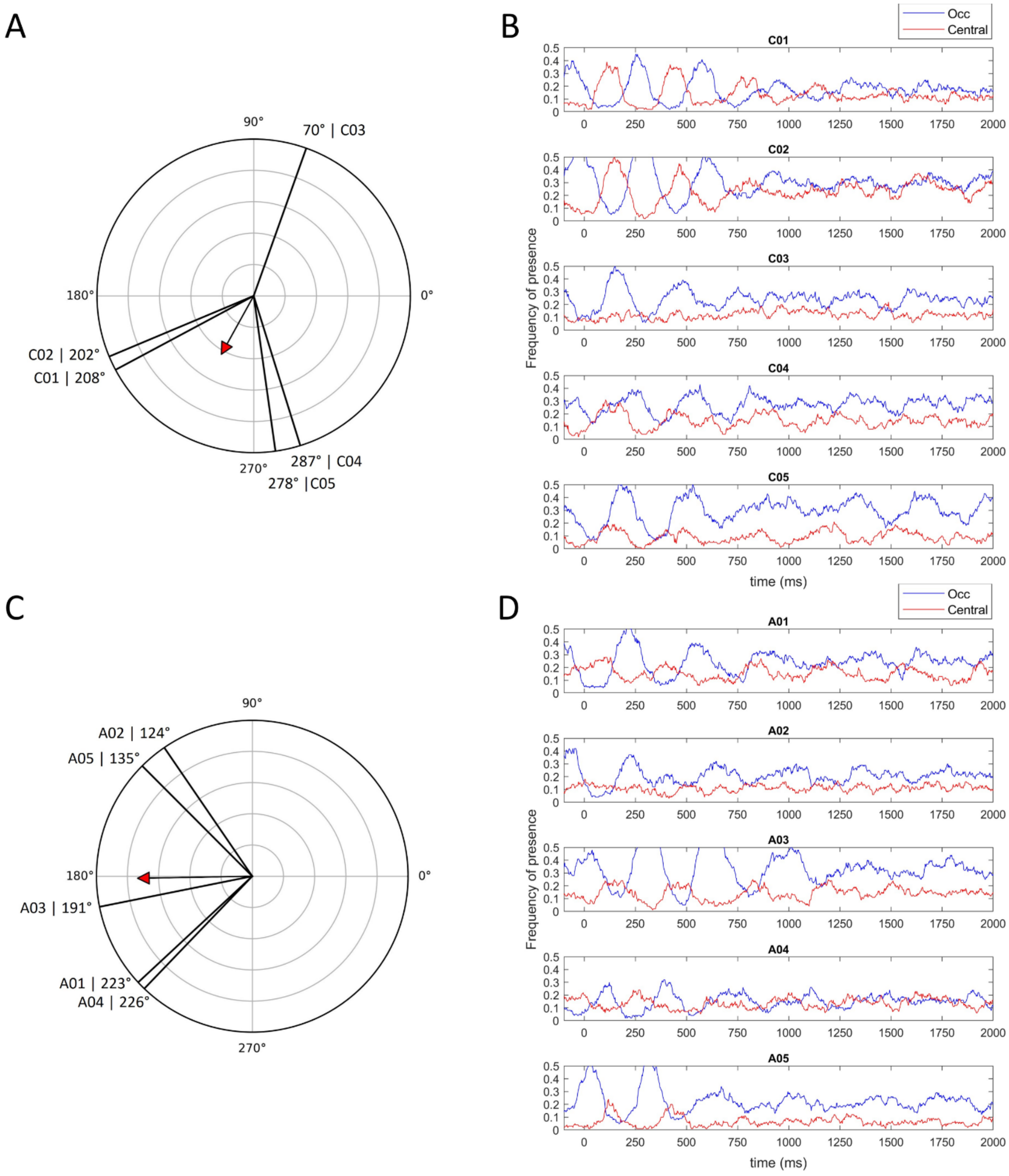
